# NeuroWRAP: integrating, validating, and sharing neurodata analysis workflows

**DOI:** 10.1101/2022.10.13.511794

**Authors:** Zac Bowen, Gudjon Magnusson, Madeline Diep, Ujjwal Ayyangar, Aleksandr Smirnov, Wolfgang Losert

## Abstract

Multiphoton calcium imaging is one of the most powerful tools in modern neuroscience. However, multiphoton data require significant pre-processing of images and post-processing of extracted signals. As a result, many algorithms and pipelines have been developed for the analysis of multiphoton data, particularly two-photon imaging data. Most current studies use one of several algorithms and pipelines that are published and publicly available, and add customized upstream and downstream analysis elements to fit the needs of individual researchers. The vast differences in algorithm choices, parameter settings, pipeline composition, and data sources combine to make collaboration difficult, and raise questions about the reproducibility and robustness of experimental results. We present our solution, called NeuroWRAP, which is a tool that wraps multiple published algorithms together, and enables integration of custom algorithms. It enables development of collaborative, shareable custom workflows and reproducible data analysis for multiphoton calcium imaging data enabling easy collaboration between researchers. NeuroWRAP implements an approach to evaluate the sensitivity and robustness of the configured pipelines. When this sensitivity analysis is applied to a crucial step of image analysis, cell segmentation, we find a substantial difference between two popular workflows, CaImAn and Suite2p. NeuroWRAP harnesses this difference by introducing consensus analysis, utilizing two workflows in conjunction to significantly increase the trustworthiness and robustness of cell segmentation results.

## Introduction

Two-photon calcium imaging is a common brain imaging technique that allows for recording the activity of hundreds or thousands of neurons at single-cell resolution. However, two-photon calcium imaging data requires significant pre-processing of acquired images (motion correction, cell segmentation, signal extraction) and post-processing of extracted signals to produce interpretable readout from each neuron. As a result, many algorithms and pipelines have been developed for the analysis of two-photon imaging data. While several algorithms and pipelines are published and available as open-source packages (Giovannucci et al., 2017; Pachitariu et al., 2017; Zhou et al., 2018; Giovannucci et al., 2019; Cantu et al., 2020), countless others are customized in individual research groups. These different analysis methods could be different ways to approach a task such as automatic cell segmentation, or algorithms may be tailored to specific use cases such as rigid versus non-rigid jitter correction. Beyond different use cases, each method or pipeline typically has a variety of input parameters such as thresholds, temporal window sizes, sampling rates, and other experimental metadata that may need to be tuned to each different dataset. The vast potential differences in algorithm choices, pipeline parameter settings, and data sources combine to create an immense challenge of analysis reproducibility and collaboration.

Reproducibility, the ability to produce the same results as previous work, is a large problem in science with neuroscience being no exception (Miłkowski et al., 2018). For the purposes of this work, we are using the word reproducibility to describe repeatability, replicability, and reproducibility. Repeatability is the ability for the same researchers to produce the same results using the same experimental setup (or software configuration) over multiple trials, which is the easiest to achieve. Replicability is the ability for different researchers to produce the same results as other researchers using the same experiment setup (or software configuration) across multiple trials. Reproducibility is the ability for different researchers to produce the same results as other researchers using a different experimental setup (or software configuration) across multiple trials (Plesser, 2018). All three are important, with replicability and reproducibility being the typical challenge point in analysis workflows causing results to differ between different researchers.

A recent study gave the same FMRI dataset to 70 different research labs with the same analysis goal and no restrictions on what analyses could be used (Botvinik-Nezer et al., 2020). All groups used different analysis pipelines and found vastly different results. The authors highlighted several solutions to this problem: 1) share less-processed data, 2) publicly share data and code 3) public pre-registration of hypothesis and analysis to be used, and 4) using consensus results from multiple pipelines. There exists the need for a robust neuroimaging data analysis platform that enforces record keeping in a way that facilitates easily reproducing past results. Efforts have been made to standardize data format in neuroscience (Teeters et al., 2015; Gorgolewski et al., 2016) as well as data storage and sharing in neuroscience (Rübel et al., 2021), but standardization of analysis techniques remains an ongoing challenge. Scientific results that can be reproduced reliably will elevate the quality of further produced work but also save trainees and early career researchers immense amounts of time when establishing data analysis workflows.

Neuroscience researchers would greatly benefit from a way to explore different options in their analysis pipelines in a controlled and reproducible manner. While analyzing neuroimaging data, it is rarely the case that the first algorithm or the first set of parameters used produce the final publishable results. It requires a significant amount of time and effort from researchers to optimize an analysis pipeline for their specific datasets because of the vast amount of parameter choices and lack of interoperability between different analysis packages due to differences in programming languages or input and output structure. Furthermore, there is not always a definitive method to know which algorithms, methods, or parameter choices are best; requiring running analyses many times and manually gathering results to determine the best configuration.

Here we present NeuroWRAP (www.neurowrap.org), a workflow integrator for reproducible analysis of two-photon data, which is an analysis platform for processing two-photon calcium imaging data. NeuroWRAP enables researchers, even with no programming experience, to analyze their data and keep an extensive record of all algorithms and parameters that were used. It facilitates reproducible analysis achieved by the ability to track, organize, and compare analysis executions, as well as allowing sharing of data and analysis workflows. Furthermore, it allows for pipelines to be easily built using algorithms from different sources and in different programming languages, such as MATLAB and Python working together seamlessly. We demonstrate how NeuroWRAP’s features aid in parameter selection in conjunction with a consensus analysis module which combines outputs of multiple algorithms to produce consistent and reliable results.

## Results

NeuroWRAP is a neurodata processing platform that contains a suite of pre-processing and analysis algorithms for two-photon calcium imaging data. It enables researchers to easily process and analyze imaging data in a reproducible and collaborative manner. NeuroWRAP focuses on analyses required to get from raw data to interpretable neuronal readout but additionally contains some downstream analysis modules. Furthermore, it is free and requires no programming experience making it accessible for all neuroscientists looking to analyze two-photon calcium imaging data. NeuroWRAP is currently available for Windows and Mac.

### Modular design

NeuroWRAP operates by executing workflows which consist of modules. Modules are the basic building blocks of workflows and consist of individual processing steps or algorithms within a typical two-photon calcium imaging analysis pipeline.

Modules are typically categorized by their processing step where the categories are as follows: 1) File acquisition, where raw image data and metadata is loaded with options for several acquisition systems such as ThorLabs (www.thorlabs.com), Bruker/Prairie (www.bruker.com), and simple TIF stacks or multipage TIFs, 2) Motion correction, which corrects motion artifacts or jitter in imaging movies, 3) Cell segmentation, where cells within the field of view are identified either automatically or manually, 4) Signal extraction, where fluorescence over time is extracted from the image stack for identified cells and baseline-corrected signals can be calculated, and finally 5) Analysis, where downstream analysis can be performed on the extracted signals. NeuroWRAP includes a library of validated modules for each of these five processing steps that can be downloaded and used (Table 1). Users of NeuroWRAP are presented with all available modules which can be individually downloaded and assigned to runtime environments as desired. The modules listed in Table 1 have been validated to work with one another following the typical dataflow from less processed (raw) data to more processed data. NeuroWRAP users can modify existing modules or write new modules from scratch. Users can then share any custom modules to other users, allowing them to see and optionally download them.

**Table 1:** Highlighted modules included in NeuroWRAP. This list is not exhaustive and will continue to expand as users share modules.

| **<u>Module Name</u>** | **<u>Type</u>** | **<u>Language</u>** | **<u>Description</u>** |
| --- | --- | --- | --- |
| Read Bruker/Prairie | File acquisition | MATLAB | Reads .raw files from Bruker/Prairie system |
| Read TIFs | File acquisition | MATLAB | Reads TIF files from a directory |
| Read Multipage TIF | File acquisition | MATLAB | Reads images from a multipage TIF file |
| Read ThorImage (Single-channel) | File acquisition | MATLAB | Reads .raw files from a ThorImage system |
| Read ThorImage (Multi-channel) | File acquisition | MATLAB | Reads .raw files from a ThorImage system with multichannel acquisition |
| Load Example Raw Data | File acquisition | MATLAB or Python | Loads an example image matrix to test workflows |
| DFT Image registration | Motion correction | MATLAB | Motion correction via Discrete Fourier Transform (DFT) registration (Guizar-Sicairos et al., 2008) |
| Suite2p Cell segmentation | Cell segmentation | Python | Automatic cell segmentation/detection from Suite2p (Pachitariu et al., 2017) |
| CaImAn cell segmentation | Cell segmentation | Python | Automatic cell segmentation/detection from CaImAn (Giovannucci et al., 2019) |
| CITE-On | Cell segmentation | Python | Automatic cell segmentation/detection from Fellin Lab |
| Manual cell selection | Cell segmentation | MATLAB | Manual cell selection via clicking GUI |
| Bandpass detection | Cell segmentation | MATLAB | Automatically detects ROIs from a mean image using bandpass filtering |
| Extract fluorescence | Signal extraction | MATLAB | Extracts fluorescence over time from detected ROIs |
| Calculate dF/F (sliding-window) | Signal extraction | MATLAB | Calculates dF/F using a sliding window to estimate baseline |
| Granger Causality Analysis | Analysis | MATLAB | Detects Granger Causal relationships between neurons based on activity (Francis et al., 2018; Francis et al., 2021) |
| Cell finder consensus | Analysis | Python | Finds matching cells from two cell segmentation algorithms |

Using modules within NeuroWRAP is significantly simpler than using packages or algorithms outside of the platform. We achieve this by requiring minimal input parameters, with default values available and visible where appropriate, and allowing outputs of modules to be piped into downstream modules as inputs. We have shared “validated” modules under the “Fraunhofer” username in NeuroWRAP (**Table 1**), which ensures module inputs and outputs are compatible with other modules in our library. As a result, any validated modules within NeuroWRAP can be used together so long as it follows the typical data flow logic of processing pipelines (i.e. from raw, less-processed data to more-processed data).

When configuring a module within a workflow, there are four different input types that can be used for input parameters:

- **<u>Manual (default)</u>:** Manual inputs let you manually enter a value into a field when creating or editing a workflow. Manual inputs support all data types except matrices.
- **<u>Runtime</u>:** Runtime inputs are just like manual inputs, but instead of entering a value while creating or editing a workflow, the value is entered prior to executing the workflow. Runtime inputs allow you to choose different input values every time you run a workflow, without editing the workflow. Runtime inputs are useful for quickly running the same analysis on many different datasets, such as when batch processing data.
- **<u>Pipe</u>:** The pipe input type enables passing data between modules by allowing you to choose an output of a prior module in the workflow as the input value. The available and compatible (i.e., matches the input’s variable type) inputs are listed in a drop-down menu. Pipe inputs allow custom workflows to be set up quickly and efficiently while also acting as a guide for data flow between modules.
- **<u>HDF5</u>:** HDF5 inputs let you choose an HDF5 file (hierarchical data format version 5; www.hdfgroup.org) and then choose a variable from that file as the input value. This input type is useful if loading pre-processed data to perform downstream analysis, or for iterating different analysis techniques on a previously executed workflow since NeuroWRAP’s output files are stored in HDF5.

When creating a workflow, all input types are viable, it is a matter of choosing which suit the user’s situation best. In most cases, early modules in a workflow will utilize mostly manual or runtime inputs, while later modules in a workflow will utilize many pipe inputs that use outputs from previous modules.

### Fully customizable workflows

With the library of available modules, workflows can be constructed to create data analysis pipelines. Workflows can be as long or short as needed, whether it is a single module or several modules. NeuroWRAP also comes with pre-built workflows which can be used as-is or easily modified to fit individual needs. All configured workflows, whether downloaded or created from scratch, will appear in the “Run Workflows” section of NeuroWRAP where they can be repeatedly executed as needed. Figure 1 show three example workflows, where the boxes between arrows represent individual modules and modules are colored according to their processing step categorization. Here we illustrate the ability to easily string together diverse algorithms, including different interpreter languages, on various types of acquired data. Additionally, one can load pre-processed data directly from an HDF5 file (utilizing the *Load from HDF5* module) or MATLAB .mat file (using the *Load from* .*mat* module) and jump right to the relevant processing step (**Fig. 1**, *bottom*). Figure 1 represents just a small fraction of the possible workflow configurations with the available module library.

**Figure 1.**
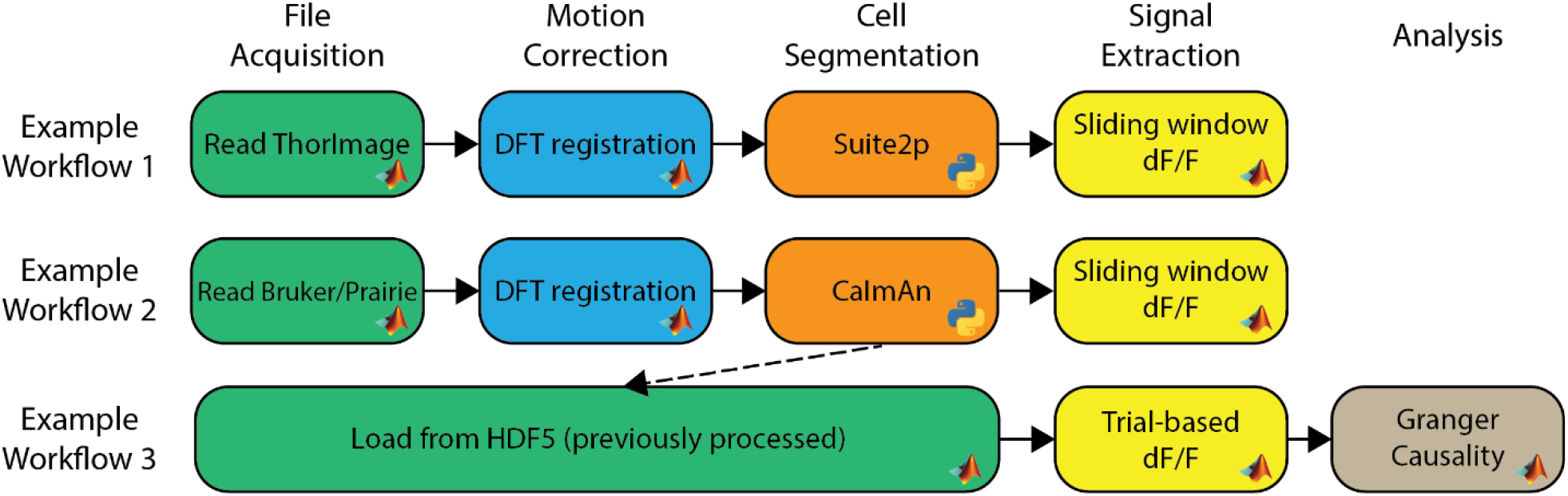
Example workflows within NeuroWRAP. These three example workflows contain modules available within NeuroWRAP. Color denotes type of processing step categorized by the column headers. Workflow 3 uses results from a previous execution to calculate dF/F using an alternative method followed by downstream analysis.

Multiple workflows can be queued up to be automatically run and each workflow will be executed in the order they are added to the queue. This workflow queue is useful if batch processing data by running the same pipeline on several different datasets or iteratively running a pipeline with altered parameters on the same dataset to explore algorithm performance. Additionally, workflows can include multiple modules from the same processing step if a comparison of results from different algorithms is desired. For example, one could place three different automatic cell segmentation modules in a workflow, which will be executed serially and then have all results available within and at the end of the workflow execution.

### Integration of algorithms and programming languages

Most existing neuroscience analysis pipelines are self-contained in that their individual algorithms are not easily mixed with algorithms from other existing analysis pipelines due to differences in programming languages, data structure, and data flow. Furthermore, most existing packages are written in either MATLAB or Python, potentially excluding users who lack the sufficient proficiency in one of these languages to be able to setup and utilize algorithms. NeuroWRAP bridges these gaps by supporting modules in both MATLAB and Python and integrating algorithms from various sources to function within a single workflow. The diversity of the module library is evident from Table 1, but it is important to note that these modules can all work together. Furthermore, modules from different interpreter language can be used seamlessly within a workflow. NeuroWRAP also includes the ability to view the module code, edit module code as needed, and write a custom module from scratch. When creating a custom module, one can specify the inputs and outputs which then generates a code template where custom procedures in python or MATLAB can be written.

NeuroWRAP features automatic handling of runtime environments to aid users that may have limited experience in setting up and managing virtual environments. When opening NeuroWRAP for the first time, a user will indicate where runtime interpreter paths are located on their computer, depending on what type of modules they wish to run. The three options for runtime environments are MATLAB, Python, and Conda. Python and Conda environments can both be used for modules written in Python, but Python runtime environments set up module requirements within a virtual environment (venv) while Conda runtime environments set up module requirements within a conda environment, a common framework for software packages in the life sciences domain. For any modules that require a Python virtual environment, the module installer automatically downloads and installs necessary dependencies based on the requirements file included with the module. MATLAB runtime environments simply point to the local path for the desired MATLAB version.

### Reproducible, recorded, and collaborative data analysis

NeuroWRAP automatically records every relevant piece of information from an analysis pipeline each time it is executed (**Fig. 2**). When a workflow is run, an execution record is created which stores all module configurations, input parameter settings, data that was parsed during execution, as well as any figures produced and resulting data. Execution records are managed within NeuroWRAP but also reside in a local directory which stores all data in an HDF5 file as well as images of any produced figures.

**Figure 2.**
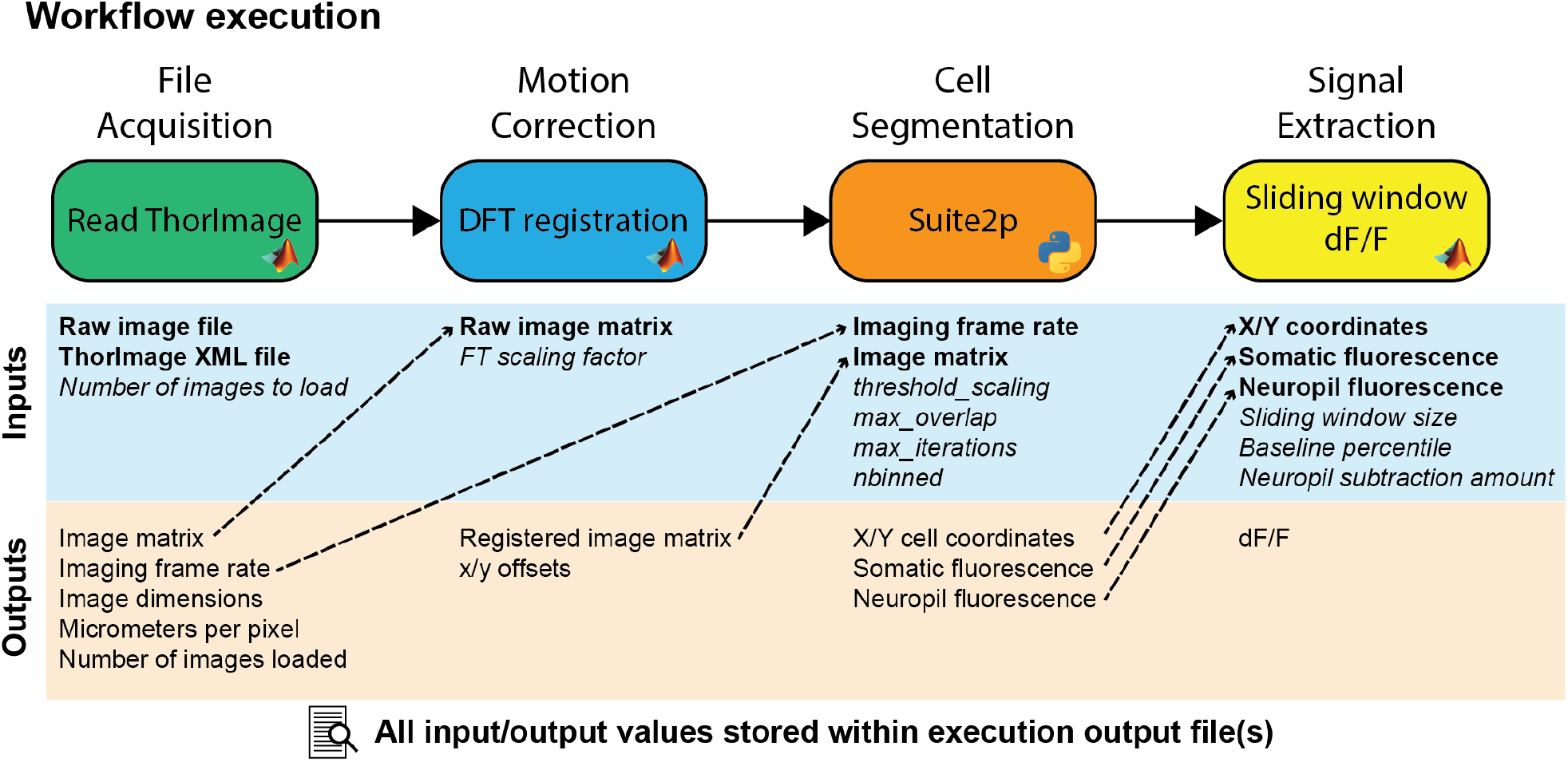
Example workflow execution. An analysis workflow featuring 4 modules. Below each module are its input and output variables, where required inputs are bolded and optional inputs are italicized. Dashed arrows represent output variables that have been piped as inputs to downstream modules. All pictured information is stored in the output file upon execution of this workflow.

Configured workflows can be shared and downloaded via NeuroWRAP enabling collaboration between users. These shared workflows can serve different purposes depending on how they are configured when shared. A shared workflow could be fully configured with all parameters preset to serve as an instructive example, or it could utilize runtime inputs so that certain key parameters can be left open for any future user. The built-in record keeping of executions facilitates reproducibility and simplifies the process of reporting analysis configuration to colleagues or in publication submissions.

### Consensus analysis of cell segmentation algorithms

Choosing an openly available algorithm to include in an analysis workflow can be difficult because even a well-configured algorithm can produce differing results from other openly available algorithms that aim to accomplish the same task. However, these differences in output of multiple algorithms can be leveraged to find which results are the most consistent and robust. To aid researchers with this challenge, we have introduced a module in NeuroWRAP that compares and combines the output of different cell segmentation algorithms, a necessary step in any calcium imaging analysis workflow. A common issue with cell segmentation algorithms that automatically detect regions of interest (ROIs) is the occurrence of false positives (non-cellular ROIs mislabeled as active neurons) and false negatives (active neurons that were not assigned ROIs). Both erroneously included ROIs or missed ROIs can negatively affect the population statistics and downstream analysis performed on the dataset. Furthermore, the occurrence of false positives and false negatives can differ between cell segmentation algorithms. An approach to solve this problem is to use the consensus of cell segmentation algorithms by utilizing detected cells that appear as accepted ROIs in both algorithms. Assuming these two unique algorithms have different types of error, consensus analysis will reveal which detected cells are the most consistent and robust because false positives are less likely to appear as a result in both algorithms. NeuroWRAP includes a cell detection consensus module which takes the output coordinates from two cell detection algorithms and returns the cells that are location-matched within a user-defined distance threshold. The distance threshold determines the maximum acceptable pixel separation of two cell locations for them to be resolved as one cell. Cells that meet the consensus criteria can then be passed to downstream modules for further processing. A NeuroWRAP workflow can be created that utilizes two separate cell detection algorithms, runs them serially, and then feeds their output to the cell detection consensus module (**Fig. 3A;** *top*). This module will also produce a visual output that shows the output of each individual cell detection module and which cells locationally-matched (**Fig. 3A;** *bottom*).

**Figure 3.**
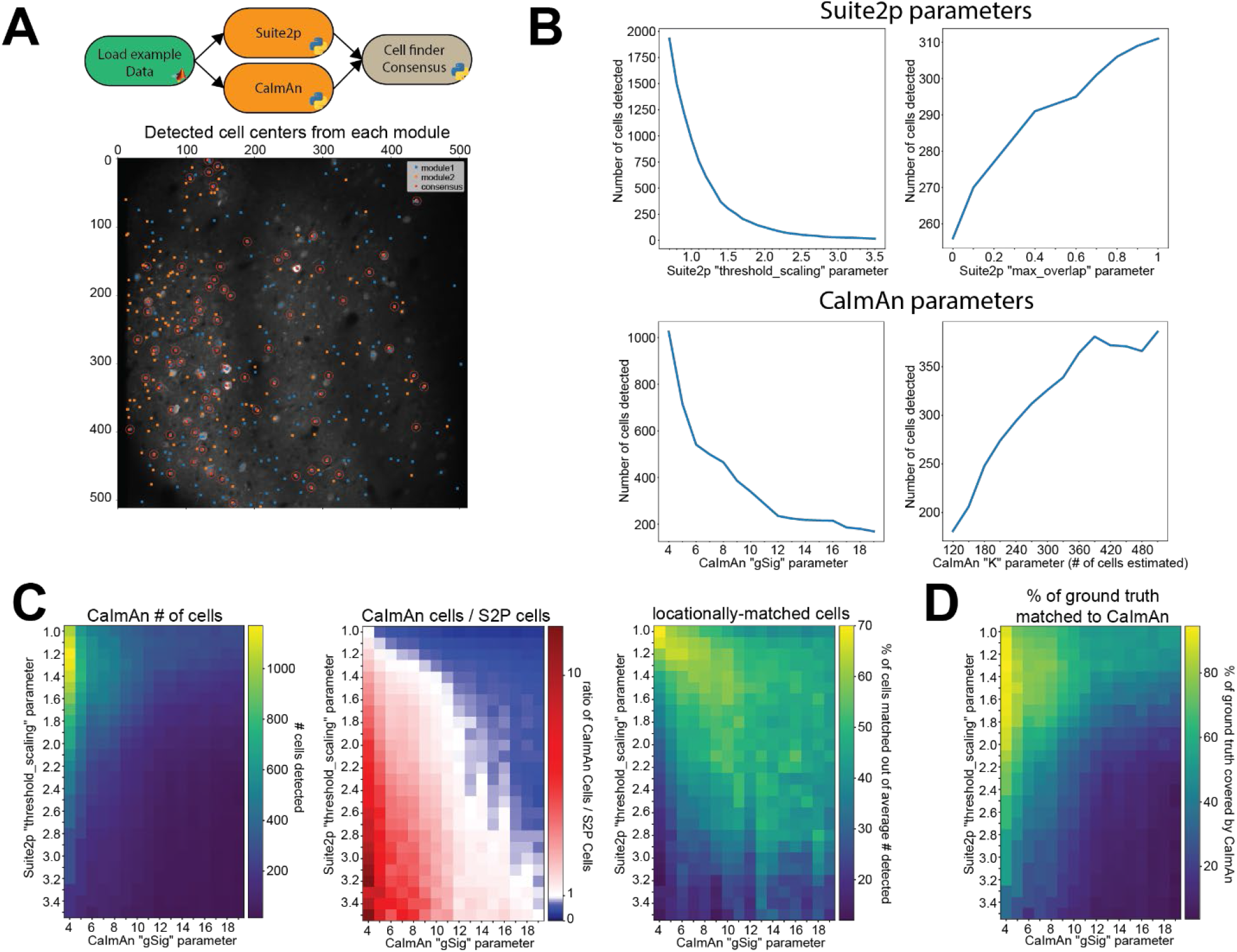
Results are sensitive to algorithm choice and parameter configuration. **A)** Example of the cell detection consensus module in NeuroWRAP. The top schematic illustrates an example usage of the Cell segmentation consensus module. The lower image is example output of the consensus module using Suite2p and CaIman on publicly available Neurofinder data. **B)** Number of cells detected by Suite2p and CaImAn while altering certain input parameters. *Top left*: “threshold_scaling”, *top right*: “max_overlap” (Suite2p), *bottom left*: “gSig” (CaImAn), *bottom right*: number of expected cells in field of view extrapolated from “K” (CaImAn). **C)** *Left*: Number of cells detected by CaImAn across a selected parameter space. *Middle*: Ratio of detected cells in CaImAn to Suite2p across a selected parameter space. White pixels indicate parameter configurations where roughly the same number of neurons were detected. *Right*: Percentage of cells that were locationally-matched between Suite2p and CaImAn at various parameter configurations. **D)** Cell-location consensus between ground truth data and CaImAn’s detected cells across a selected parameter space.

### NeuroWRAP enables crucial parameter exploration

An overarching concern when considering results from neuroscience data analysis is that due to various reasons, researchers may not explore many different analysis procedures and settings on all of their data. This may be due to difficulty in tracking results across many different configurations, lacking of understanding of how sensitive parameters may be, or simply a lack of time. As a result, researchers may use parameter settings based on what was historically used in their lab or parameter settings used in well-cited literature, regardless of whether these settings are appropriate for their newly acquired data. Furthermore, slight differences in parameter choices can produce drastically different results, leading to a propagation of variability with downstream analyses and ultimately different answers to the scientific question being tested. Effort spent exploring and recording results from different algorithms and parameter settings could lead to more robust and reproducible science.

The flexibility of modular workflows in NeuroWRAP is designed to enable users to easily compare different algorithms and parameter choices when constructing an analysis pipeline. To demonstrate this and further motivate the value of exploring different modules and different parameter settings within analysis pipelines, we constructed example workflows that are identical up to the point of cell segmentation, where they differ. We use automatic cell segmentation algorithms from two popular imaging analysis packages, Suite2p (Pachitariu et al., 2017) and CaImAn (Giovannucci et al., 2019). These workflows were run on the same publicly available dataset which has a ground truth of 330 cells determined from anatomical labelling (See Materials and Methods). To start, we view how the results from these modules change when sensitive parameters are adjusted. For Suite2p, we tested the number of cells detected as we altered two parameters: the *threshold_scaling* parameter which controls the threshold requirement for the signal-to-noise ratio on each ROI, and the *max_overlap* parameter which determines the amount of overlap two ROIs can have to be deemed unique. We found that the *threshold_scaling* parameter drastically affects the number of cells detected (**Fig. 3B**; *top left*) in an expected trend with lower values significantly overestimating the cell count and higher values underestimating the cell count. The *max_overlap* parameter followed an expected trend as well, yet with changes that were less drastic in magnitude (**Fig. 3B**; *top right*). For CaImAn, we again tested the number of cells detected as we altered two parameters: the *gSig* parameter which is the expected half-size of cells and *K* which is the number of expected cells per patch (approximated from total estimate in the entire field of view). We found that the *gSig* parameter has a large effect on the number of cells detected, with lower, unrealistic values drastically overestimating the number of cells, with mid-range values producing values closer to ground truth, though still with variation (**Fig. 3B**; *bottom left*). The *K* parameter had more surprising results, with certain values causing CaImAn to find more cells than estimated, while other values lead CaImAn to find fewer cells than estimated (**Fig. 3B;** *bottom right*).

These examples of parameter exploration highlight the sensitivity of workflow configuration when it comes to producing consistent results. A simple exploration of a reduced range of these values could help users make informed decisions on parameter selections. For example, if a researcher knows the size of the field of view and the average cell size, an approximation can be made about how many cells are expected in the field of view. Utilizing NeuroWRAP to iteratively analyze data at a range of parameter values to explore how many cells are found in a recorded and reproducible manner can aid researchers in choosing an appropriate parameter setting.

### Consensus analysis can aid in parameter configuration of modules

We have shown how individual parameters can drastically alter results and need to be carefully considering when setting up analysis pipelines. We next explore how this information can be further utilized by comparing and combining the outputs of both modules and using the consensus analysis of detected cells, utilizing the unique advantages and features of NeuroWRAP. We test how these two example cell detection algorithms, Suite2p and CaImAn, agree or deviate from one another under various parameter configurations. To make the results comparison as close as possible, we use the number of cells found by Suite2p to seed CaImAn’s expected number of cells detected (parameter *K* from the previous section). We choose to alter the two most sensitive parameters from the previous section: *threshold*_scaling within Suite2p and *gSig* within CaImAn. We find that the number of cells detected by CaImAn expectedly decrease with higher threshold_scaling and gSig values, following the trend found in each individual algorithm (**Fig. 3C;** *left*). However, with the ratio of number of cells detected by CaImAn to number of cells detected by Suite2p (**Fig. 3C**; *middle*), we find that there is a narrow region of the parameter space (diagonally from top left to bottom right) where the two algorithms are finding roughly the same number of cells. This narrow region of the parameter space where two different algorithms agree gives more confidence in how parameters should be configured for robust results. However, the number of cells on its own may not be fully informative since the cell locations may differ. Therefore, we utilize the consensus module and determine which neurons are spatially-matched at each joint parameter configuration (**Fig. 3C;** *right*). We find a similar region of high consensus between the two algorithms indicating how parameters in these two algorithms can be tuned to produce consistent output.

Together these results point towards favorable regions within the parameter space where different algorithms reach high consensus, and parameter selection can be made according to regions of high consensus (**Fig. 3C**; *middle, right*) and whether the researcher wants more or fewer cells detected (**Fig. 3C**; *left*) depending on their research context. For example, if studying which cells are highly responsive across many trials in a stimulus-based experimental paradigm, one may wish to pick a portion of this favorable region where fewer cells are being detected (toward the bottom right), assuming highly active cells take precedent in these particular cell detection algorithms. Conversely, if studying population dynamics at a fine temporal scale, one may utilize a portion of this region to capture the activity of as many cells as possible (towards the top left) to capture the joint activity of all possible neurons. In either case, using a reduced and more coarsely sampled range of parameters in conjunction with consensus analysis could serve as a useful tool when determining how best to configure analysis on one’s data. This process is made significantly easier within the context of NeuroWRAP where all configurations are recorded with each workflow execution.

Utilizing the ground truth data from anatomical labelling, we next compare the cell locations found by CaImAn across the parameter space to assess their spatial accuracy (**Fig. 3D**) as follows:

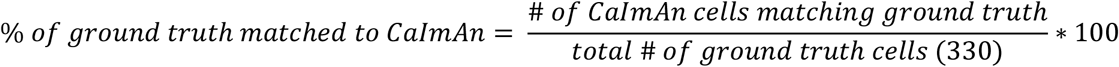

We find a bias such that the more cells that are detected by CaImAn, the more likely there are to be matches with the ground truth data. Given that there is finite space within the field of view and a restriction on overlap of cells, the regions of high percentage matched with the ground truth data should still be taken as favorable to those regions with a low percentage. However, in the pursuit of proper parameter selection for cell detection, ground truth labelling is typically absent and thus consensus analysis may hold more merit.

Lastly, we utilize the ground truth data to determine whether consensus cells are more likely to be real and therefore more robust. We first look at how many of CaImAn’s detected cells are spatially-matched with ground truth at each parameter value (**Fig. 4A**) as follows:

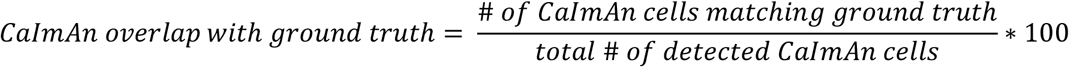

**Figure 4.**
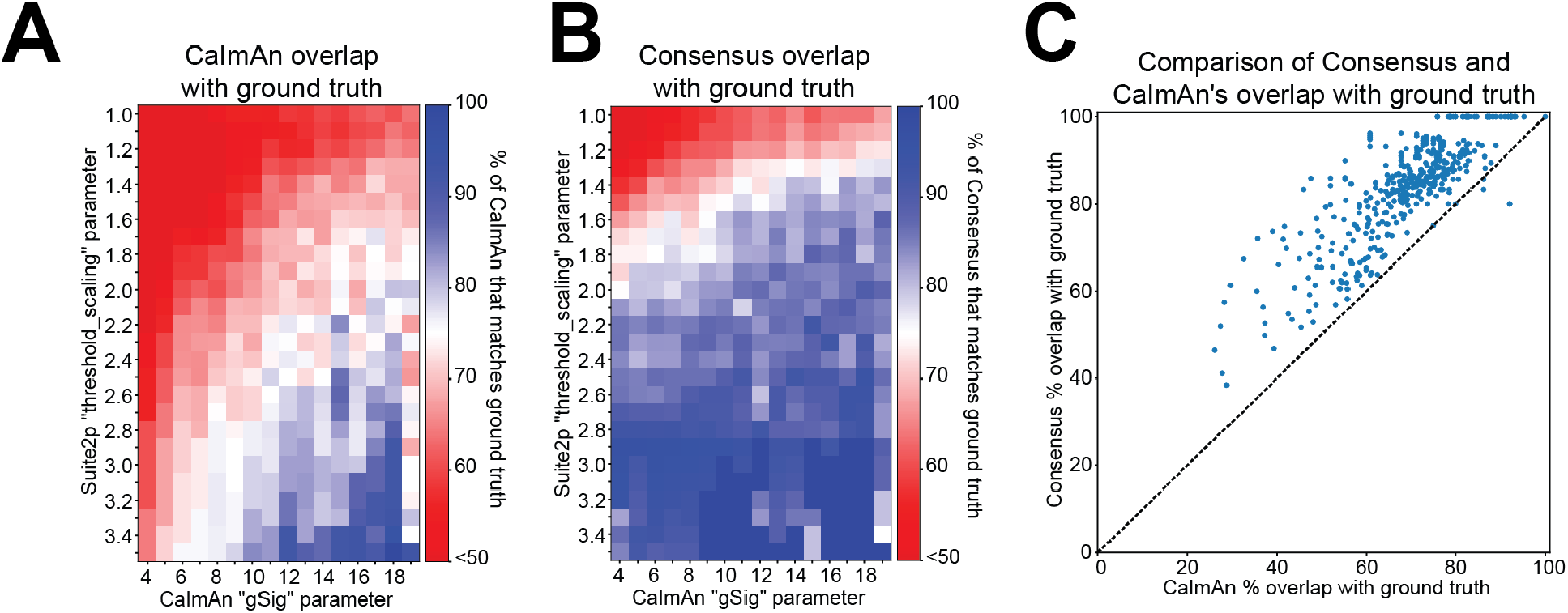
Cell detection comparisons to ground truth data. **A)** Proportion of CaImAn’s detected cells that overlap with ground truth cell locations at a range of parameter configurations. **B)** Proportion of Consensus analysis (Suite2p and CaImAn) cell locations that overlap with ground truth cell locations at a range of parameter configurations. **C)** Scatter plot of each pixel value in plots **A** and **B**, comparing the percentage overlap with ground truth from CaImAn alone to that of CaImAn’s consensus with Suite2p.

We find that for most of the parameter space, only 75% or less of CaImAn’s cells are spatially-matched with ground truth. It’s important to note that CaImAn is tailored towards detecting active cells, and some of the ground truth data may contain inactive cells, which may account for some of the discrepancy. Additionally, the extremes of this parameter space may be drastically overestimating or underestimating the number of cells, accounting for further deviations from ground truth. We next look at what proportion of the consensus cells, meaning the spatially-matched cells that CaImAn and Suite2p both found, also overlap with ground truth cell locations (**Fig. 4B**) as follows:

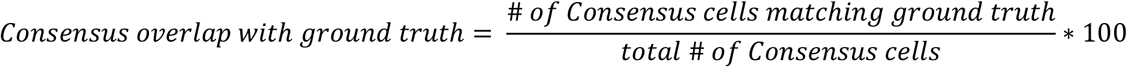

We see that consensus cell locations have a high overlap with ground truth (greater than 75%) at a wide range of the parameter space. We next compare the ground truth overlap of CaImAn and consensus cell locations by plotting the percent overlap with ground truth of each against one another at each parameter configuration (**Fig. 4C**). We see that most points lie above the diagonal, indicating that consensus has a higher proportion of its cells spatially-matched with ground truth, regardless of parameter configuration.

Consensus analysis allows researchers to be more lenient with parameter selection since the agreement cells are more likely to real and robust, while outliers or false positives would have to be produced by both individual algorithms to exist in the consensus results. In a real analysis scenario, ground truth data will likely be unavailable, but these results indicate that using the consensus cell locations rather than one cell detection algorithm in isolation awards greater confidence in the detected cell locations, especially when exploring several different parameter configurations.

## Discussion

Here we have presented NeuroWRAP, a workflow integrator for reproducible analysis of two-photon data. NeuroWRAP allows researchers to process and analyze multiphoton calcium imaging data while allowing them to explore, record, and share every aspect of their analysis pipeline. NeuroWRAP provides an analysis environment that contains a suite of options for each processing step in a calcium imaging data analysis pipeline while promoting reproducibility and collaboration between researchers. Furthermore, NeuroWRAP encourages researchers to explore ways to test their data analysis workflows and make them more robust. We have motivated this point with consensus analysis and parameter selection (**Fig. 3**).

Many options exist for analysis of calcium imaging data, both in the form of individual algorithms that handle a single processing step, to suites that handle the full analysis pipeline. NeuroWRAP does not aim to replace existing algorithms, but rather aggregate and wrap existing tools in single environment where they can work together. For example, Suite2p (Pachitariu et al., 2017) and CaImAn (Giovannucci et al., 2019) are two popular options that each contain all of the analysis needed to process calcium imaging data. However, these tools are not intended to be modularized and used interchangeably with other algorithms. NeuroWRAP accomplishes interoperability between these algorithms as well as many more (see Table 1).

As the tools available for processing multiphoton imaging data continue to develop and be extended, NeuroWRAP is designed to enable future extension seamlessly. Users of NeuroWRAP can extend its library and capabilities simply by sharing modules that they create within NeuroWRAP or sharing workflows that they construct using existing modules. Our team will continue to incorporate new algorithms and pipelines into NeuroWRAP as they are popularized or requested.

Comparative and consensus analysis aim to address the challenge of variabilities in the analysis pipelines and results. Comparative and consensus analysis (i) investigates how comparable analysis should be compared with one another, (ii) generates insights to how one result differs or is similar to another, and (iii) offers mechanisms to consolidate the results and generate a consensus in order to produce a more reliable/trustworthy result. We have demonstrated that simple consensus based on agreement of two algorithms is sufficient to significantly enhance the robustness of the cell segmentation pipeline to parameter choice, and to enhance the trustworthiness of the selected cells using ground truth experimental data. Ultimately, the consensus analysis leads us to recommend choosing cell segmentation parameters that slightly overestimated the number of cells and pruning based on consensus.

Note that the application of comparative and consensus analysis can occur in any stage in the experimental pipeline; for example, performing such analysis on the pre-processing pipeline can result in a more robust downstream analysis. The concept of consensus analysis has been proposed in software engineering domain, including in machine learning. The N-version programming technique (i.e., multiple versions of functionally equivalent programs are independently developed from the same software specification) along with majority voting algorithm has been used in safety-critical software systems to make decisions. In machine learning, multiple predictors can be used to produce the final estimate or predictor – these are often referred to “agreement of experts” or “ensemble learning”, and various consensus rules exist.

Future work on NeuroWRAP will focus on further integrating consensus analysis across the workflow and making the platform applicable for a broader range of research data, in particular incorporating more experimental metadata such as behavioral readout and trial classification which will provide the necessary experimental information to incorporate more downstream analysis modules as well.

## Materials and Methods

### Consensus analysis test data

Data used for figure production (Figure 3) was downloaded from the publicly available datasets as part of the Neurofinder challenge (http://neurofinder.codeneuro.org/). Figure 3 and Figure 4 utilize dataset N00.00 from the Svoboda lab and was acquired from an awake head-fixed mouse expressing genetically encoded calcium indicator GCaMP6s. Ground truth labelled ROIs were used for comparison to detected cell locations, where the precise x-and y-coordinate of each ground truth ROI was calculated as the average of the ROI pixel locations.

### Cell-finder consensus analysis

The cell-finder consensus module uses two sets of input cell coordinates and finds which points are within the user-defined pixel distance threshold. When two cell coordinates are within this distance threshold, the average of the two cell locations is taken as the consensus coordinate location. For the figures in this work, a pixel distance of 15 was used, meaning cell centers within 15 pixels (roughly one cell diameter) are merged into one.

### Parameter exploration analysis

For Suite2p analysis, parameters were kept close to default values except when altered for parameter exploration or to fit the characteristics of the data (such as image sampling rate). Parameter “threshold_scaling” was altered between 0.7 and 3.4 in 0.1 increments while max_overlap was held at the default value of 0.75. During alterations of the max_overlap value between 0 and 1 in 0.1 increments, the threshold_scaling parameter was held at 1.5.

For CaImAn analysis, parameters *K, rf*, and *stride* were set according to the CaImAn documentation to appropriately estimate the cell density. *gSig* is the expected half-size of neurons in pixels (approximate neuronal radius). Parameter *rf* is the half-size of patches and was set to *gSig*\*4, while parameter *stride* is the overlap between patches in pixels and was set to *gSig*\*2. Parameter *K* is the expected number of components per patch which we computed as *K = K_total/npatches* where *npatches* was determined by patch size and *K_total* was set according to estimate number of components in the entire field of view. For gSig analysis in Figure 3B, we set *K_total=330 neurons* as it was the number of cells labelled in the ground truth dataset. For all other analysis, *K_total* was varied across a pre-defined range (**Fig. 3B**; *bottom right*) or according to the number of cells detected by Suite2p (**Fig. 3C**,**D**).

## Notes

**Conflicts of interest:** None

### Competing Interest Statement

The authors have declared no competing interest.

